# Direct RNA sequencing reveals SARS-CoV-2 m6A sites and possible differential DRACH motif methylation among variants

**DOI:** 10.1101/2021.08.24.457397

**Authors:** João H. Campos, Juliana T. Maricato, Carla T. Braconi, Fernando Antoneli, Luiz Mario R. Janini, Marcelo R. S. Briones

## Abstract

The causative agent of COVID-19 pandemic, the SARS-CoV-2 coronavirus, has a 29,903 bases positive-sense single-stranded RNA genome. RNAs exhibit about 100 modified bases that are essential for proper function. Among internal modified bases, the *N^6^*-methyladenosine, or m6A, is the most frequent, and is implicated in SARS-CoV-2 immune response evasion. Although the SARS-CoV-2 genome is RNA, almost all genomes sequenced so far are in fact, reverse transcribed complementary DNAs. This process reduces the true complexity of these viral genomes because incorporation of dNTPs hides RNA base modifications. Here, in this perspective paper, we present an initial exploration of the Nanopore direct RNA sequencing to assess the m6A residues in the SARS-CoV-2 sequences of ORF3a, E, M, ORF6, ORF7a, ORF7b, ORF8, N, ORF10 and the 3’-untranslated region. We identified 15 m6A methylated positions, of which, 6 are in ORF N. Also, because m6A is associated with the DRACH motif, we compared its distribution in major SARS-CoV-2 variants. Although DRACH is highly conserved among variants we show that variants Beta and Eta have a fourth position C>U change in DRACH at 28,884b that could affect methylation. The Nanopore technology offers a unique opportunity for the study of viral epitranscriptomics. This technique is PCR-free and is not sequencing-by-synthesis, therefore, no PCR bias and synthesis errors are introduced. The modified bases are preserved and assessed directly with no need for chemical treatments or antibodies. This is the first report of direct RNA sequencing of a Brazilian SARS-CoV-2 sample coupled with the identification of modified bases.

## INTRODUCTION

RNA viruses are causative agents of major human transmissible diseases such as influenza, poliomyelitis, measles and COVID-19 (Hogle, 2002; Guerra et al., 2017; Krammer et al., 2018; Islam et al., 2021; Wang et al., 2021). In the Baltimore classification, viruses with RNA genomes comprise Groups III, IV, V and VI while DNA viruses are in Groups I, II and VII (Baltimore, 1971; Dill et al., 2016). COVID-19 is a highly contagious viral disease with severe respiratory, inflammatory and thrombotic manifestations (Hanff et al., 2020; Jiang et al., 2020). The COVID-19 pandemic is caused by a Beta coronavirus, SARS-CoV-2, included in Group IV of the Baltimore classification (Almeida et al., 1968; Cui et al., 2019). In SARS-CoV-2 the RNAs serve as information storage when packaged into the viral particle and as mRNAs for viral protein synthesis upon infection of mammalian cells (Wu et al., 2020; V’kovski et al., 2021).

The SARS-CoV-2 genome consists of a positive-sense single-stranded strand RNA with 29,903 bases (V’kovski et al., 2021). There are approximately 100 different base modifications in all RNA species, and these modified bases are essential for proper translation, splicing and RNA metabolism (Brocard et al., 2017). Among these modified bases, the *N^6^*-methyladenosine (m6A) is the most frequent internal base modification, and is found in viruses with exclusive cytoplasm replication, such as Zika Virus, Dengue virus and Hepatitis C virus (Gokhale et al., 2016). Methylated base m6A is implicated in SARS-CoV-2 evasion of the host immune response because the methylated viral RNA does not bind to the host protein RIG-I (retinoic acid-inducible gene I), responsible for the type-1 interferon (IFN1) response, an activator of immune pathways (Brocard et al., 2017; Li et al., 2021). The viral RNA is m6A methylated by host’s methylases METTL3, METTL14, WATAP and KIAA1429, called “writers”, and demethylated by FTO and ALKBH5, called “erasers” which normally methylate the host’s RNAs (Gatsiou and Stellos, 2018). Knockdown of METTL3 significantly reduces the SARS-CoV-2 methylation and blocks the viral mechanism of RIG-I binding inhibition (Li et al., 2021).

Almost all SARS-CoV-2 genomes sequenced so far are reverse transcribed complementary DNAs (cDNAs) although the genome is, in fact, RNA (Nazario-Toole et al., 2021). Reverse transcription provides a fast, practical, PCR prone method for sequencing the SARS-CoV-2 genome. However, it reduces the true complexity of these viral genomes. The incorporation of dNTPs in the first strand cDNA chain makes the RNA base modifications, present in the RNA template chain, mostly indistinguishable from unmodified bases and sequencing errors (Kietrys et al., 2017). RNA modified bases are critical for proper biological function and are involved in several diseases, encompassing the field of Epitranscriptomics Medicine (Gatsiou and Stellos, 2018). Although several different technologies have been used for identification of modified bases in mRNAs and viral RNAs they require large quantities of material which precludes single-cell analysis and low abundance samples. Also, the antibody-dependent methylation analysis does not provide nucleotide level accuracy of modified bases (Brocard et al., 2017; Jenjaroenpun et al., 2021; Li et al., 2021).

The Oxford Nanopore Technology (ONT) has been used for SARS-CoV-2 whole-genome cDNA sequencing (Bull et al., 2020). Also, this same technology has been used for direct RNA sequencing of SARS-CoV-2 (Kim et al., 2020; Taiaroa et al., 2020; Vacca et al., 2020). The major advantage of ONT direct RNA sequencing over cDNA sequencing is the identification of modified bases. Two previous studies on SARS-CoV-2 direct RNA sequencing did not couple the genetic analysis with base modification identification while a third study detected the 5mC methylation using this technology (Kim et al., 2020; Taiaroa et al., 2020; Vacca et al., 2020).

In this perspective paper we assessed the potential of the Nanopore direct RNA sequencing for identification of m6A residues, at nucleotide level resolution, in the SARS-CoV-2 genome. For this we analyzed direct RNA sequencing reads of open reading frames (ORFs) 3a, E, M, 6, 7a, 7b, 8, N, 10 and the 3’-untranslated region (Jenjaroenpun et al., 2021). In addition, since m6A is associated with the DRACH motif (D = G/A/U, R = G/A, H = A/U/C), we compared the DRACH distribution in major SARS-CoV-2 variants to verify if potential variant-specific alterations in m6A methylation patterns occur in SARS-CoV-2 evolution (Bayoumi and Munir, 2021).

## MATERIAL AND METHODS

All procedures for viral isolation and initial passages were performed in a biosafety level 3 laboratory (BLS3), in accordance with WHO recommendations and under the laboratory biosafety guidance required for the SARS-CoV-2 at the BLS3 facilities at the Federal University of São Paulo. SARS-CoV-2 stock was kindly provided by Prof. José Luiz Proença-Módena (University of Campinas – UNICAMP, SP, Brazil).

### SARS-CoV-2 infections

For SARS-CoV-2 infection, the Vero E6 cell line (ATCC® CRL-1586™) was maintained in Minimum Essential Medium (MEM; Gibco) supplemented with 10% fetal bovine serum (FBS) (Gibco) and 1% penicillin/streptomycin (Gibco). Vero E6 cells were kept in a humidified 37°C incubator with 5% CO2. After reaching 80% confluent monolayer, cells were seeded in 24-well plates at a density of 5 × 10^5^ cells per well. Cells were infected at 1 × 10^5^ PFU/well (MOI 0.2) with SARS-CoV-2 lineage B (Araujo et al., 2020), 3^rd^ passage and kept for 2 hours at 37°C with 5% CO2 in MEM supplemented with 2.5% FBS and 1% penicillin/streptomycin. Cells were rinsed with 1x PBS to remove attached viral particles and fresh MEM with 10% FBS was added to the cultures. After 48 hours the cell cultures were halted and used for supernatant harvesting (Hoffmann et al., 2020).

### RNA Extraction

RNA samples from culture supernatants were extracted using viral QIAamp Viral RNA Mini Kit (Qiagen, USA). Briefly, 350 μL of supernatants were centrifuged at 3,000 rpm for 5 minutes to remove cell debris and transferred to new tubes containing 550 μL of lysis buffer (AVL - provided with the kit) and RNA isolation was performed according to the manufacturer’s instructions. RNA samples were quantitated with Nanodrop (Thermo Fischer Scientific, USA).

### Direct RNA sequencing

For RNA sequencing, 9.5μl of RNA containing ≈50ng of RNA, from Vero E6 cells supernatant, were used in the Nanopore RNA Sequencing SQK-RNA002 following the manufacturer’s instructions (Nanopore Technologies). Runs were performed in MinION (Oxford Nanopore) with flowcell FLO-MIN106 for 40 hours and 1.59 million reads were generated. Raw data (fast5 files) were used for basecalling with Guppy (v-5.0.11), in high-accuracy mode.

### Assembly

The resulting fastq reads were aligned to the SARS-CoV-2 reference (GISAID ID: EPI_ISL_413016) with minimap2 (v-2.21-r1071) (Li, 2018). The resulting sam files were converted to bam files and all reads were sorted and indexed according to the reference coordinates using samtools (v-1.13) (Li et al., 2009). The “index” and “eventalign” modules of nanopolish (v-0.13.3) were used to generate an index of base call reads for the signals measured by the sequencer, and to align events to the reference transcriptome, checking for differences in current that may suggest modifications in the base.

### Methylation analysis

The probability of methylation in DRACH motifs was calculated with m6anet (v-0.1.1-pre) (Hendra and Wan, 2021), as recommended: (I) to preprocess the segmented raw signal file with “m6anet-dataprep”, and (II) run m6anet over data using “m6anet-run_inference”.

### DRACH motif comparison

Comparative analysis and annotation of DRACH motifs (Bayoumi and Munir, 2021), identified with m6Anet (v-01.1-pre) (Hendra and Wan, 2021), among SARS-CoV-2 variants were carried out using Geneious v-10.4 (http://www.geneious.com). Five sequences of each variant isolated and sequenced in Brazil were aligned using the Geneious aligner. For variant Eta only one sequence from Brazil is available from GISAID (http://www.gisaid.org) that is complete with high coverage and therefore samples from the US, France, Spain, and Canada were used. The SARS-CoV-2 variants sequences analyzed are deposited in the GISAID database (http://www.gisaid.org) with accession numbers for Alpha (EPI_ISL_1133259, 1133268, 1133267, 1495029, 3316204), Beta (1966629, 1742275, 1966124, 1445171, 1716877), Gamma (3539883, 3539773, 3545813, 3545803, 3540000), Delta (3540039, 3540020, 3540001, 3460250, 3505224), Eta (1583653, 3502618, 3535614, 3490821, 3493947), Lambda (1445272, 3010903, 2928137, 1966094, 2617911) and, Zeta (3434818, 2841610, 3506974, 3190295, 1494963).

## RESULTS

The assembly of the 3’-half of the SARS-CoV-2 RNA genome was obtained by mapping the RNA reads to GISAID ID: EPI_ISL_413016, the strain used for infection of Vero E6 cells (**Figure 1A**). The coverage varied from 30x to 1,600x from 5’ to 3’ starting at ORF3a. Because the RNA sequencing adapter is ligated to the 3’-ends of RNAs the coverage is higher as it gets closer to the 3’-end. Several reads reached the “Spike” ORF but were not used for further analysis due to low coverage. Sequencing runs of 40 hours were sufficient to obtain around 2,000 reads corresponding to the 3’-half of the SARS-CoV-2 genome. The average sequence length was 787 bases and the mode 1,350 bases. The global identity to the reference was 90%. The phred score of the assembled bases are between 20 and 30. ORF N has a substantial coverage because of its proximity to the 3’-end (≈1,000x).

**FIGURE 1.**
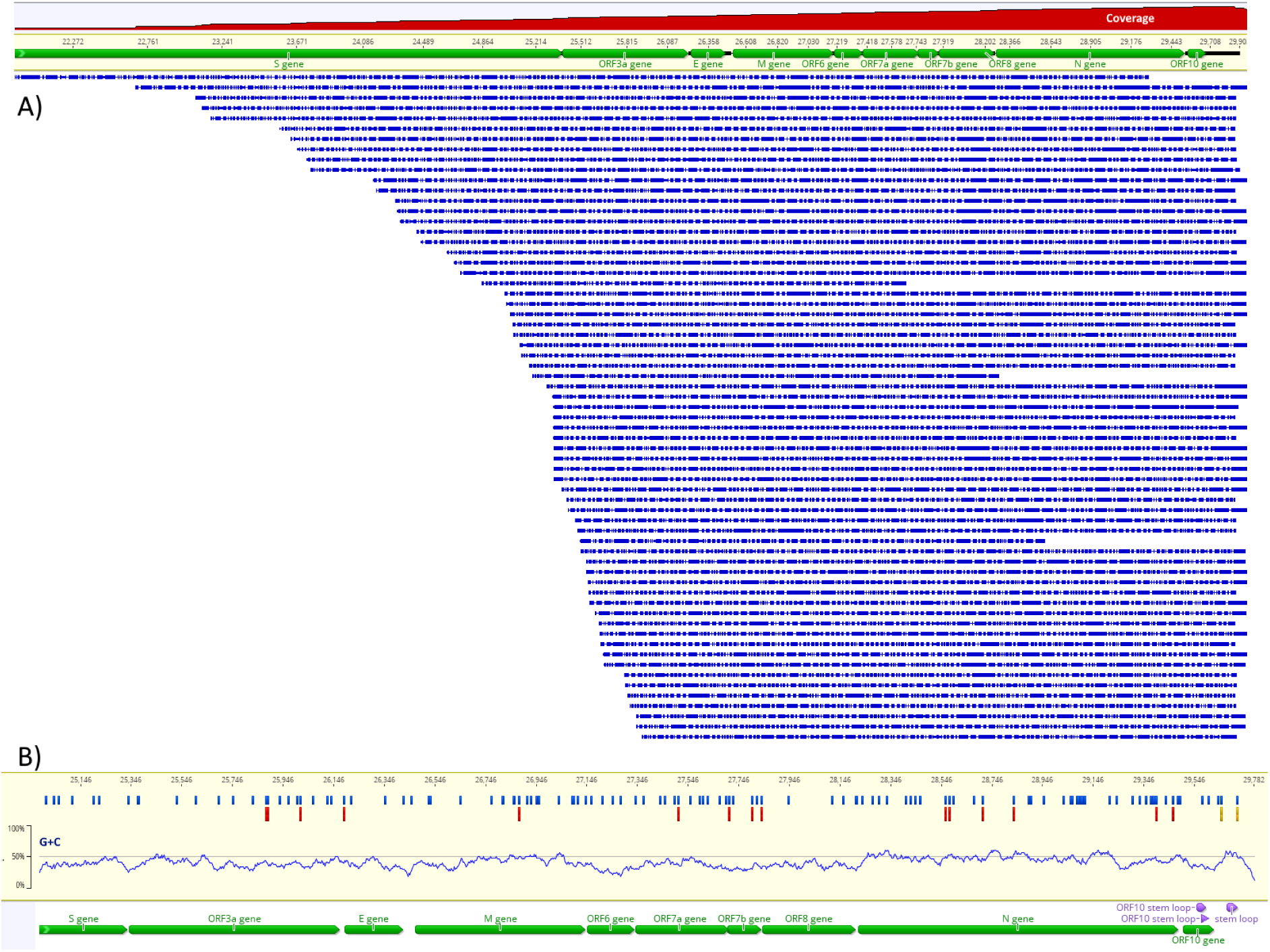
Assembly of Nanopore direct RNA reads to the reference sequence (A) and map of m6A methylation probability along the SARS-CoV-2 RNA (B). In (A) green horizontal bars indicate the ORFs, blue horizontal bars of decreasing size indicate the Nanopore reads and the red area at the top indicates the log scale coverage from 1x to 1,600x. In (B) blue vertical bars indicate the DRACH motifs, red vertical bars indicate m6A (>50% probability), the yellow vertical bars indicate two potential m6A with probabilities 0.38 and 0.44 in the 3’-untranslated region. The G+C content is indicated in the plot just above the annotation.

Direct RNA sequencing reads were used for detection of m6A using the m6Anet tool, validated by systematic benchmark (Chen et al., 2021). This method uses the reference sequence, the basecalled fastq files and the raw fast5 files to identify the DRACH motifs and the raw signal data in fast5 files with corresponding signal alterations associated with m6A to calculate the probability of bona fide methylation (Chen et al., 2021). Using this approach, we identified 15 positions within DRACH with >50% methylation probability (**Figure 1B**). Among these positions, 11 have more than 100x coverage and 4 positions have >80% methylation probability (**Table 1**). The nucleocapsid region (N) was the most enriched methylated region with six m6A residues and at least 30 DRACH motifs.

**TABLE 1.**
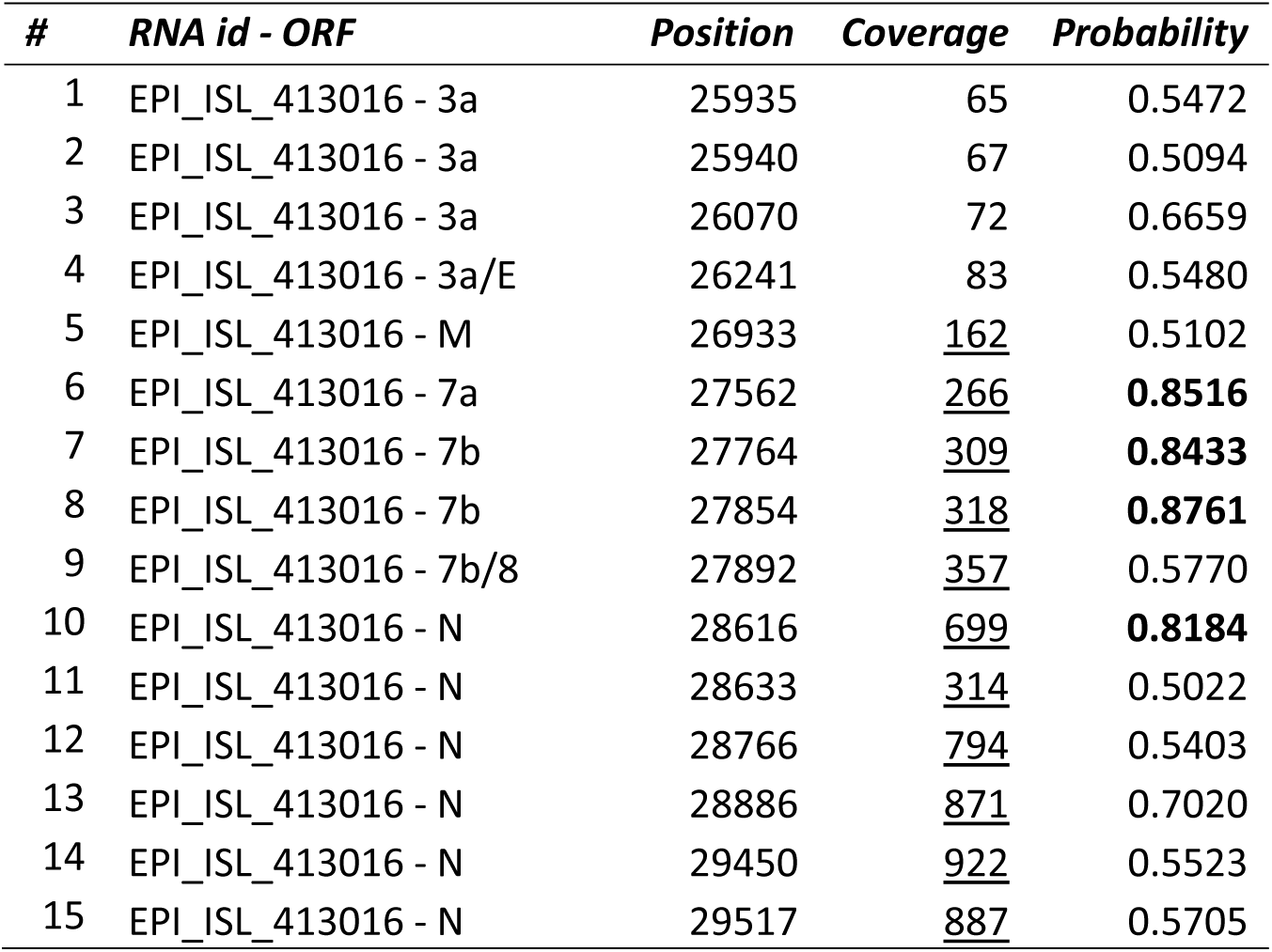
Distribution and sequencing coverage of potential methylated adenosines in SARS-CoV-2 RNA genome. Coverage > 100x is underlined and probability > 80 is in boldface as calculated by m6Anet (Hendra and Wan, 2021). After m6Anet analysis only sites with coverage above 60 were considered. Position numbering according to Wuhan reference sequence (GenBank NC_045512).

| # | RNA id - ORF | Position | Coverage | Probability |
| --- | --- | --- | --- | --- |
| 1 | EPI_ISL_413016 - 3a | 25935 | 65 | 0.5472 |
| 2 | EPI_ISL_413016 - 3a | 25940 | 67 | 0.5094 |
| 3 | EPI_ISL_413016 - 3a | 26070 | 72 | 0.6659 |
| 4 | EPI_ISL_413016 - 3a/E | 26241 | 83 | 0.5480 |
| 5 | EPI_ISL_413016 - M | 26933 | <u>162</u> | 0.5102 |
| 6 | EPI_ISL_413016 - 7a | 27562 | <u>266</u> | **0.8516** |
| 7 | EPI_ISL_413016 - 7b | 27764 | <u>309</u> | **0.8433** |
| 8 | EPI_ISL_413016 - 7b | 27854 | <u>318</u> | **0.8761** |
| 9 | EPI_ISL_413016 - 7b/8 | 27892 | <u>357</u> | 0.5770 |
| 10 | EPI_ISL_413016 - N | 28616 | <u>699</u> | **0.8184** |
| 11 | EPI_ISL_413016 - N | 28633 | <u>314</u> | 0.5022 |
| 12 | EPI_ISL_413016 - N | 28766 | <u>794</u> | 0.5403 |
| 13 | EPI_ISL_413016 - N | 28886 | <u>871</u> | 0.7020 |
| 14 | EPI_ISL_413016 - N | 29450 | <u>922</u> | 0.5523 |
| 15 | EPI_ISL_413016 - N | 29517 | <u>887</u> | 0.5705 |

Because the DRACH motif is associated with m6A we tested if the major SARS-CoV-2 variants, Alpha, Beta, Gamma, Delta, Eta, Lambda and Zeta isolated in Brazil exhibited mutations within DRACH that differ between variants. The alignment consisted of the Wuhan reference sequence (GenBank NC_045512), the Brazilian reference (GISAID ID: EPI_ISL_413016) and five sequences of major variants (Material and Methods). DRACH is highly conserved among variants, however, differences can be observed (**Figure 2**). Five sequences of the variant Beta and 4 of the variant Eta have a C>U mutation in the fourth position of DRACH (at position 28,886) that could block methylation at this site (**Figure 2B, D1**). The methylation probability at this site is 70% and the coverage 871x. Another change in DRACH that could probably interfere with methylation is a C>U at 28,947 (**Figure 2B, D2**) in a single variant Zeta sequence, although the methylation probability at this site is <50%. Other DRACH variants observed are probably “silent” such as the 4 nucleotides insertion in the intergenic region between ORF8 and ORF N in the five sequences of variant Gamma (**Figure 2B, D3**). The insertion does not change the DRACH sequence.

**FIGURE 2.**
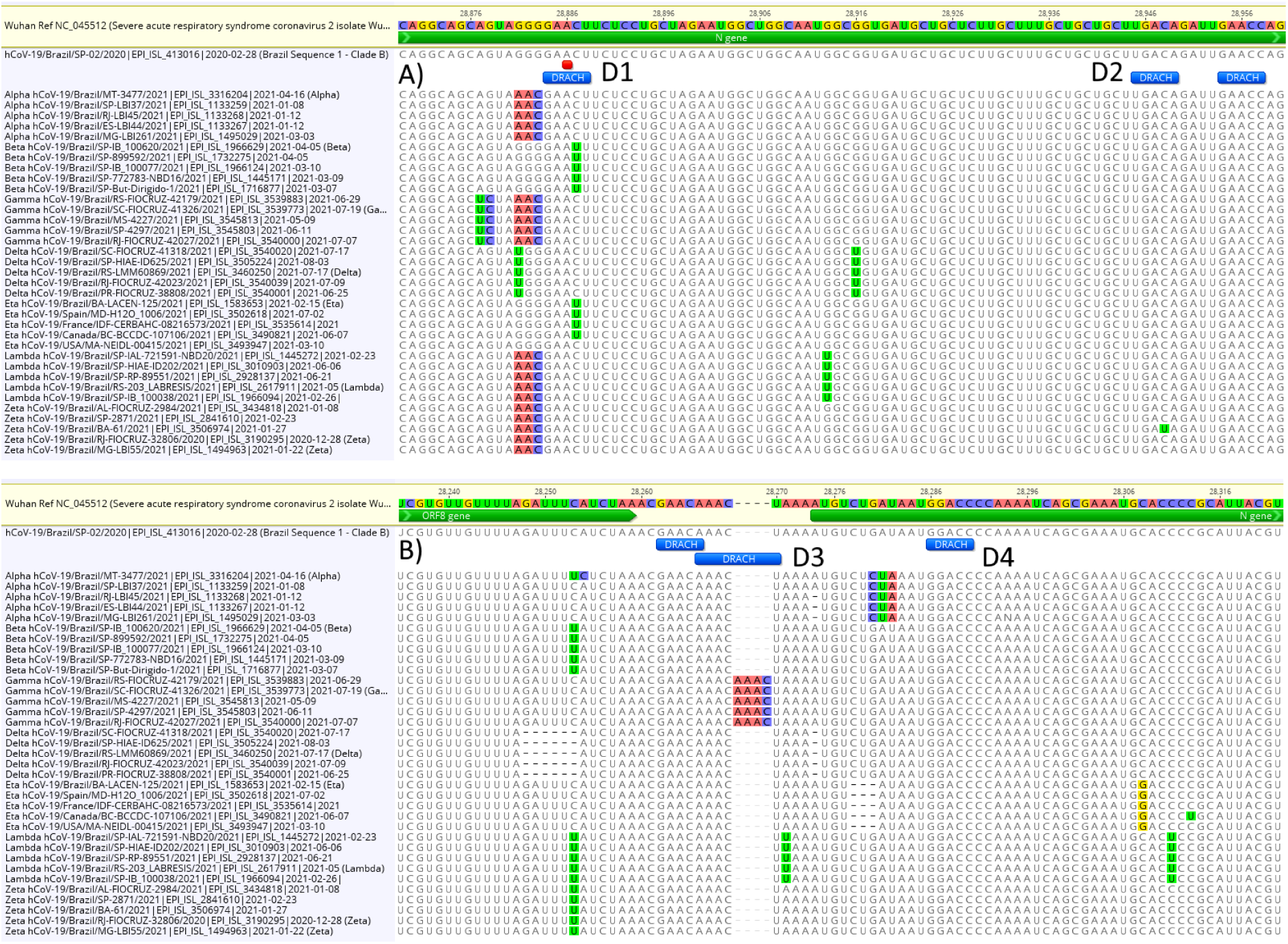
Variability of DRACH sequences among SARS-CoV-2 variants. In (A) the C>U mutation at 28,886 disrupts the m6A (marked red) DRACH motif D1 in five variant Beta sequences and four variant Eta sequences while a single variant Zeta has a C>U mutation that disrupts the DRACH motif D2. In (B) a 4 bases insertion in the intergenic region ORF8/N does not disrupt DRACH D3 while D4 shows a fully conserved DRACH.

## DISCUSSION

RNA modification, or epitranscriptomics, is a key factor in viral infections (Brocard et al., 2017). Methylation of RNA bases, either as host’s multitargets or viral RNAs are involved in immunity and associated disease progression (Li et al., 2021, 3; Meng et al., 2021). Various types of RNA methylation are involved in Covid-19 immunity and novel proposed treatments for COVID-19 involve methylase inhibitors (Ahmed-Belkacem et al., 2020; Paramasivam, 2020). The Nanopore direct RNA sequencing offers a unique opportunity for the study of viral epitranscriptomics (Kim et al., 2020; Jenjaroenpun et al., 2021). Direct RNA sequencing is PCR-free, is not sequencing-by-synthesis and, therefore, not affected by PCR bias and synthesis errors. The modified bases are preserved and assessed directly with no need for chemical treatments or antibodies. Direct RNA sequencing of SARS-CoV-2 has been validated by orthogonal methods (Vacca et al., 2020).

In the present study we show that direct RNA sequencing of SARS-CoV-2 is an important tool for the assessment of the full complexity of this viral RNA genome. Identification of m6A was performed in supernatants of SARS-CoV-2 infected Vero E6 cells and, therefore, is enriched in genomic RNAs and not subgenomic RNA. Sequencing was performed without PCR amplification or any other *in vitro* synthesis (**Figure 1**). Mapping of m6A using direct RNA sequencing data was achieved by comparison of raw and basecalled reads and the m6A pattern identified was fully consistent with the nucleocapsid region m6A enrichment observed by liquid chromatography-tandem mass spectrometry and methylated RNA immunoprecipitation sequencing (MeRIP-seq) (Li et al., 2021). The probability of m6A, as calculated by the m6Anet method, was >50% and a minimum coverage of 60x in four positions and the recommended 100x coverage in 11 positions (Chen et al., 2021). Our results show that the nucleocapsid region is the most heavily methylated (**Table 1**), confirming previous studies by Met-RIP, mass spectrometry (for m6A) and direct RNA sequencing (for 5mC) (Kim et al., 2020; Li et al., 2021).

As expected, the DRACH pattern is highly conserved among SARS-CoV-2 variants although in, at least one position, a significant mutation was observed in the nucleocapsid region, in variants Beta and Eta (**Figure 2**) that could negatively affect methylation by disrupting the DRACH motif (Bayoumi and Munir, 2021). The significance of this finding needs to be further investigated with comparative infection experiments to determine the impact of this mutation in viral growth of these variants.

Although direct RNA has been used for SARS-CoV-2 sequencing, none of these studies coupled the sequencing with base modification analysis, the major advantage of direct RNA sequencing (Taiaroa et al., 2020; Vacca et al., 2020). The study that couples SARS-CoV-2 direct RNA sequencing with methylation analysis, identified 5mC but not m6A. This present work is the first report of direct RNA sequencing, coupled with identification of modified m6A bases, of a Brazilian SARS-CoV-2 isolate.

As a future perspective we are working on a methodology to allow full length, high coverage, sequencing of the whole SARS-CoV-2 RNA genome by direct RNA sequencing and therefore extend the m6A analysis to ORF1ab, Spike and the 5’-untranslated region.

## ACKNOWLEDGEMENTS

The authors thank Prof. José Luiz Proença-Módena (University of Campinas – UNICAMP, SP, Brazil) for providing the SARS-CoV-2 sample and Fernanda Prieto, M.Sc., (Interprise, Inc.) for suggestions and assistance with Nanopore supply acquisitions.

## DATA AVAILABILITY STATEMENT

The datasets presented in this study will be available upon request.

## AUTHOR CONTRIBUTIONS

JTM and CTB maintained cell lines and viruses, collected cells and purified RNA. MRSB prepared the sequencing libraries, performed the RNA sequencing, basecalling and DRACH analysis and wrote the initial manuscript. JHC performed bioinformatics analysis, basecalling, sequence assembly and modified bases analysis. FA assisted with bioinformatics analysis. LMJ assisted with experimental design and data analysis. All authors discussed the results and edited the manuscript.

## REFERENCES

Ahmed-Belkacem, R., Sutto-Ortiz, P., Guiraud, M., Canard, B., Vasseur, J.-J., Decroly, E., et al. (2020). Synthesis of adenine dinucleosides SAM analogs as specific inhibitors of SARS-CoV nsp14 RNA cap guanine-N7-methyltransferase. Eur J Med Chem 201, 112557. doi:10.1016/j.ejmech.2020.112557.

Almeida, J. D., Berry, D. M., Cunningham, C. H., Hamre, D., Hofstad, M. S., Mallucci, L., et al. (1968). Virology: Coronaviruses. Nature 220, 650–650. doi:10.1038/220650b0.

Araujo, D. B., Machado, R. R. G., Amgarten, D. E., Malta, F. de M., de Araujo, G. G., Monteiro, C. O., et al. (2020). SARS-CoV-2 isolation from the first reported patients in Brazil and establishment of a coordinated task network. Mem Inst Oswaldo Cruz 115, e200342. doi:10.1590/0074-02760200342.

Baltimore, D. (1971). Expression of animal virus genomes. Bacteriol Rev 35, 235–241. doi:10.1128/br.35.3.235-241.1971.

Bayoumi, M., and Munir, M. (2021). Evolutionary conservation of the DRACH signatures of potential N6-methyladenosine (m6A) sites among influenza A viruses. Sci Rep 11, 4548. doi:10.1038/s41598-021-84007-0.

Brocard, M., Ruggieri, A., and Locker, N. (2017). m6A RNA methylation, a new hallmark in virus-host interactions. J Gen Virol 98, 2207–2214. doi:10.1099/jgv.0.000910.

Bull, R. A., Adikari, T. N., Ferguson, J. M., Hammond, J. M., Stevanovski, I., Beukers, A. G., et al. (2020). Analytical validity of nanopore sequencing for rapid SARS-CoV-2 genome analysis. Nat Commun 11, 6272. doi:10.1038/s41467-020-20075-6.

Chen, Y., Davidson, N. M., Wan, Y. K., Patel, H., Yao, F., Low, H. M., et al. (2021). A systematic benchmark of Nanopore long read RNA sequencing for transcript level analysis in human cell lines. bioRxiv, 2021.04.21.440736. doi:10.1101/2021.04.21.440736.

Cui, J., Li, F., and Shi, Z.-L. (2019). Origin and evolution of pathogenic coronaviruses. Nat Rev Microbiol 17, 181–192. doi:10.1038/s41579-018-0118-9.

Dill, J. A., Camus, A. C., Leary, J. H., Di Giallonardo, F., Holmes, E. C., and Ng, T. F. F. (2016). Distinct Viral Lineages from Fish and Amphibians Reveal the Complex Evolutionary History of Hepadnaviruses. J Virol 90, 7920–7933. doi:10.1128/JVI.00832-16.

Gatsiou, A., and Stellos, K. (2018). Dawn of Epitranscriptomic Medicine. Circ Genom Precis Med 11, e001927. doi:10.1161/CIRCGEN.118.001927.

Gokhale, N. S., McIntyre, A. B. R., McFadden, M. J., Roder, A. E., Kennedy, E. M., Gandara, J. A., et al. (2016). N6-Methyladenosine in Flaviviridae Viral RNA Genomes Regulates Infection. Cell Host & Microbe 20, 654–665. doi:10.1016/j.chom.2016.09.015.

Guerra, F. M., Bolotin, S., Lim, G., Heffernan, J., Deeks, S. L., Li, Y., et al. (2017). The basic reproduction number (R0) of measles: a systematic review. The Lancet Infectious Diseases 17, e420–e428. doi:10.1016/S1473-3099(17)30307-9.

Hanff, T. C., Mohareb, A. M., Giri, J., Cohen, J. B., and Chirinos, J. A. (2020). Thrombosis in COVID-19. Am J Hematol 95, 1578–1589. doi:10.1002/ajh.25982.

Hendra, C., and Wan, Y. K. (2021). GoekeLab/m6anet: Pre-release. Zenodo doi:10.5281/zenodo.4692776.

Hoffmann, M., Kleine-weber, H., Schroeder, S., Krüger, N., Herrler, T., Erichsen, S., et al. (2020). SARS-CoV-2 Cell Entry Depends on ACE2 and TMPRSS2 and Is Blocked by a Clinically Proven Protease Inhibitor. Cell 181, 271–280.

Hogle, J. M. (2002). Poliovirus Cell Entry: Common Structural Themes in Viral Cell Entry Pathways. Annu Rev Microbiol 56, 677–702. doi:10.1146/annurev.micro.56.012302.160757.

Islam, M. A., Kundu, S., Alam, S. S., Hossan, T., Kamal, M. A., and Hassan, R. (2021). Prevalence and characteristics of fever in adult and paediatric patients with coronavirus disease 2019 (COVID-19): A systematic review and meta-analysis of 17515 patients. PLoS One 16, e0249788. doi:10.1371/journal.pone.0249788.

Jenjaroenpun, P., Wongsurawat, T., Wadley, T. D., Wassenaar, T. M., Liu, J., Dai, Q., et al. (2021). Decoding the epitranscriptional landscape from native RNA sequences. Nucleic Acids Res 49, e7. doi:10.1093/nar/gkaa620.

Jiang, S., Xia, S., Ying, T., and Lu, L. (2020). A novel coronavirus (2019-nCoV) causing pneumonia-associated respiratory syndrome. Cell Mol Immunol 17, 554–554. doi:10.1038/s41423-020-0372-4.

Kietrys, A. M., Velema, W. A., and Kool, E. T. (2017). Fingerprints of Modified RNA Bases from Deep Sequencing Profiles. J Am Chem Soc 139, 17074–17081. doi:10.1021/jacs.7b07914.

Kim, D., Lee, J.-Y., Yang, J.-S., Kim, J. W., Kim, V. N., and Chang, H. (2020). The Architecture of SARS-CoV-2 Transcriptome. Cell 181, 914–921.e10. doi:10.1016/j.cell.2020.04.011.

Krammer, F., Smith, G. J. D., Fouchier, R. A. M., Peiris, M., Kedzierska, K., Doherty, P. C., et al. (2018). Influenza. Nat Rev Dis Primers 4, 1–21. doi:10.1038/s41572-018-0002-y.

Li, H. (2018). Minimap2: pairwise alignment for nucleotide sequences. Bioinformatics 34, 3094–3100. doi:10.1093/bioinformatics/bty191.

Li, H., Handsaker, B., Wysoker, A., Fennell, T., Ruan, J., Homer, N., et al. (2009). The Sequence Alignment/Map format and SAMtools. Bioinformatics 25, 2078–2079. doi:10.1093/bioinformatics/btp352.

Li, N., Hui, H., Bray, B., Gonzalez, G. M., Zeller, M., Anderson, K. G., et al. (2021). METTL3 regulates viral m6A RNA modification and host cell innate immune responses during SARS-CoV-2 infection. Cell Rep 35, 109091. doi:10.1016/j.celrep.2021.109091.

Meng, Y., Zhang, Q., Wang, K., Zhang, X., Yang, R., Bi, K., et al. (2021). RBM15-mediated N6-methyladenosine modification affects COVID-19 severity by regulating the expression of multitarget genes. Cell Death Dis 12, 1–10. doi:10.1038/s41419-021-04012-z.

Nazario-Toole, A. E., Xia, H., and Gibbons, T. F. (2021). Whole-genome Sequencing of SARS-CoV-2: Using Phylogeny and Structural Modeling to Contextualize Local Viral Evolution. Mil Med, usab031. doi:10.1093/milmed/usab031.

Paramasivam, A. (2020). RNA 2’-O-methylation modification and its implication in COVID-19 immunity. Cell Death Discov 6, 118. doi:10.1038/s41420-020-00358-z.

Taiaroa, G., Rawlinson, D., Featherstone, L., Pitt, M., Caly, L., Druce, J., et al. (2020). Direct RNA sequencing and early evolution of SARS-CoV-2. doi:10.1101/2020.03.05.976167.

Vacca, D., Fiannaca, A., Tramuto, F., Cancila, V., Paglia, L. L., Mazzucco, W., et al. (2020). Direct RNA nanopore sequencing of SARS-CoV-2 extracted from critical material from swabs. doi:10.1101/2020.12.21.20191346.

V’kovski, P., Kratzel, A., Steiner, S., Stalder, H., and Thiel, V. (2021). Coronavirus biology and replication: implications for SARS-CoV-2. Nat Rev Microbiol 19, 155–170. doi:10.1038/s41579-020-00468-6.

Wang, C., Wang, Z., Wang, G., Lau, J. Y.-N., Zhang, K., and Li, W. (2021). COVID-19 in early 2021: current status and looking forward. Sig Transduct Target Ther 6, 1–14. doi:10.1038/s41392-021-00527-1.

Wu, C., Liu, Y., Yang, Y., Zhang, P., Zhong, W., Wang, Y., et al. (2020). Analysis of therapeutic targets for SARS-CoV-2 and discovery of potential drugs by computational methods. Acta Pharmaceutica Sinica B 10, 766–788. doi:10.1016/j.apsb.2020.02.008.

